# Heterochrony of puberty in the European Badger (*Meles meles*) can be explained by growth rate and group-size: Evidence for two endocrinological phenotypes

**DOI:** 10.1101/405803

**Authors:** Nadine Adrianna Sugianto, Chris Newman, David Whyte Macdonald, Christina Dagmar Buesching

## Abstract

Puberty is a key stage in mammalian ontogeny, involving endocrinological, physiological and behavioural changes, moderated by intrinsic and extrinsic factors. Thus, not all individuals within one population achieve sexual maturity simultaneously. Here, using the European badger (*Meles meles*) as a model, we describe male testosterone and female oestrone profiles (using Enzyme-immunoassays) from first capture (3 months, post-weaning) until 28 months (attaining sexual maturity and final body size), along with metrics of somatic growth, scent gland development and maturation of external reproductive organs as well as intra-specific competition. In both sexes, endocrinological puberty commenced at ca. 11 months. Thereafter, cub hormone levels followed adult seasonal hormone patterns but at lower levels, with the majority of cubs reaching sexual maturity during their second mating season (22-28 months). Interestingly, there was evidence for two endocrinological phenotypes among male cubs (less evident in females), with early developers reaching sexual maturity at 11 months (first mating season) and late developers reaching sexual maturity at 22-26 months (second mating season). Early developers also attained a greater proportion of their ultimate adult size by 11 months, exhibiting faster growth rates than late developers (despite having similar adult size). Male cubs born into larger social groups tended to follow the late developer phenotype. Our results support the hypothesis that a minimum body size is required to reach sexual maturity, which may be achieved at different ages, even within a single population, where early maturity can confer individual fitness advantages and enhance population growth rate.

## Introduction

Puberty represents a key stage in mammalian ontogeny during which a variety of endocrinological, physiological, and behavioural changes occur [1]. It is marked by the development of secondary sexual characteristics [2], the first occurence of ovulation/oestrus in females and the onset of spermatogenesis in males [3]. During puberty, typically both sexual and somatic maturation are completed [4], but some species may continue to grow even after reaching sexual maturity [5]. Although the age at puberty depends predominantly on intrinsic genetic factors, its timing can be moderated by a variety of additional extrinsic factors, such as food availability, seasonal variation, environmental conditions [6-8] and/ or the presence of conspecifics [3], as well as dynamic interactions between these factors [4]. Thus, typically not all members of a species [9], or even all individuals within one population [1, 10-13] mature simultaneously or at the same rate, leading to heterochrony [14-15]. In mammals, the onset of puberty typically depends on attaining a minimum body size (often a certain functional proportion of the final adult body size: [10,16], and conspecifics may reach this minimum required body size at different ages (e.g., dairy calves require 56-60% of adult body weight which they may reach between 49.8 and 58.2 weeks of age: [10]). Individuals that experience restricted resources during development therefore tend to undergo puberty at an older age than do individuals that experienced abundant resourses and are thus in better nutritional condition [13], consequently there can be a trade-off between somatic growth and puberty/ reproductive activity [17].

A distinctive endocrinological feature of puberty is the full activation of the hypothalamic-pituitary-gonadal (HPG) axis. This involves the episodic release of gonadotropin releasing hormones (GnRH) by the hyphothalamus, which in turn activates the anterior pituitary gland to secrete luteinizing hormone (LH) and follicle stimulating hormone (FSH) that instigate the generation of gamets and release of sex steroids [18]: in males, LH stimulates testosterone production from the interstitial cells of the testes (Leydig cells), and FSH stimulates testicular growth and enhances the production of an androgen-binding protein by the Sertoli cells, which are a component of the testicular tubules necessary for sustaining maturing sperm cells [2]. In females, FSH stimulates the ovarian follicle(s), causing one/ several ovum/ ova to grow, and triggers the production of follicular oestrogen. This rise in oestrogen then causes the pituitary gland to cease production of FSH and to increase LH production instead. This rise in LH levels in turn causes the ovum/ ova to be released from the ovary, resulting in ovulation [2]. Therefore, oestrogen and testosterone levels are low throughout the prepubertal period, but increase immediately prior, during and after puberty, until they reach adult concentrations [19-20].

Nevertheless, although the physiological processes of puberty have been the subject of numerous studies [6, 8, 21], for many mammals, especially in wild-living populations, knowledge regarding the factors driving sexual development and potential heterochrony in puberty onset is still lacking [7, 20-21]. Particularly in seasonal breeders, it is often difficult to determine sexual maturity unambiguously, because even adults exhibit periods of reproductive quiescence when males cease spermatogenesis and females do not undergo oestrus cycles [22-23]. Most carnivores (for exceptions see [24-26]), and all mustelids [27], undergo such periods of seasonal reproductive quiescence, and in many the timing of their sexual maturity is still being evaluated [26]. Here, we use the European badger (*Meles meles*; henceforth “badger”) as a model seasonally breeding carnivore to investigate the endocrinological changes and concomitant ontological development of male and female genitalia during puberty, and examine how onset of puberty can be affected by body size and intra-specific competition resulting in developmental heterochrony.

### Badger reproduction and development

The badgers’ mating season is restricted mainly to January-March, although further matings can occur throughout the summer months [28] with local population density determining the number of additional oestrous cycles ranging from nil to monthly [29-30]. During the mating season, scent marking activity increases [31], where particularly the subcaudal gland secretion plays an important role in group-cohesion and olfactory mate guarding [32-33], as well as resource defense and reproductive advertisement [34-35]. This is reflected by significant elevation in the production of subcaudal secretion [32] as well as changes to the secretion’s chemical composition [34]. During the mating season, all mature males have large scrotal testes, and females exhibit a distinctly swollen, pink and everted vulva [36]. In contrast, during autumnal reproductive quiescence, males have smaller testes that ascend into the body cavity, while females cease to exhibit vulval swelling [36]. Sex steroid levels also exhibit distinct seasonal patterns in sexually mature badgers [37, 30, 36]: In males, testosterone levels are high in spring and summer, low in autumn and peak during the winter mating season. In females, oestrone levels are high in spring, low in summer, peak in autumn and remain elevated for pregnant females in winter but decline in non-pregnant females. As in all carnivores, male badgers have a bacculum (os penis; [38]) that provides mechanical support during copulation, thus enabling prolonged intromission and mate-guarding through copulatory tying [39-40], facilitates sperm transport [41], and helps trigger ovulation in species with induced ovulation such as badgers [28].

Badgers produce one litter annually (with mean litter size at our study site = 1.4±0.06, range of 1-4 cubs, where 93% litters comprise less than 3 cubs: [42]), born between mid January – mid March (76% of cubs in the UK are born in mid-February: [42]). Newborn cubs are highly altricial, and their eyes and ear canals do not open until they reach 5 weeks of age [43]. Cubs are weaned at 6-8 weeks of age [44], during which time they first emerge from their underground den, termed a sett [45], and are fully integrated into the social group at 14-16 weeks of age [45]. Growth rate has been shown to vary among cubs depending on resource availability, which is affected by prevailing weather conditions, linked to food (mainly earthworm) availability [46] and natal sett quality [47-48], as well as infection with a highly pathologic intestinal coccidian (*Eimeria melis*), where surviving infected male and female cubs exhibit, respectively, 5 cm and 3.5 cm shorter mature body-length [49].

In high density areas (such as our study population in Wytham Woods: 44.55 ± 5.37 (SE) badgers/km^2^; [42, 50]), cubs take longer to reach adult size than in lower density areas [51, 52] and remain smaller than those living at lower density [51, 53]). Cubs start producing subcaudal gland secretion when they are approximately 4 months old [32] but anoint themselves with secretion from adults (a behaviour termed ‘scent-theft’) at a much younger age [45], signifying the importance of this secretion in badger sociality [33]. Reports of sexual maturity vary considerably, ranging from 9-12 months [54-55] to 18 months [32]; but most studies evade the issue of exact age at puberty and simply state that female badgers would not be able to breed until reaching the age of 2 years due to delayed embryonic implantation (reviewed in [43]). No studies to-date have investigated potential developmental heterochrony between individuals within the same population or cub cohort.

Here, we describe for the first time male testosterone and female oestrone profiles for cubs, commencing from the time of first capture (i.e., at the end of the closed season when cubs are 3 months of age and are fully weaned) through to the age of 28 months (i.e., when all badgers have reached sexual maturity: [54]; as well as full adult size: [52]), and report on the ontological development of male and female external genitalia morphology (EGM; i.e., degree of testicular descent and vulval swelling), bacculum length, and testes volume. Because in other mammals, somatic growth as well as sexual maturity have been shown to vary among individuals within the same population [1,10], we then investigate if all cubs in our sample mature at the same rate, or if there is evidence for different ontological strategies in terms of hormone profiles, skeletal growth and the production of subcaudal gland secretion (as reported in other species: [16]).

## Materials and methods

### Badger Trapping and Sampling

Data were collected from a high-density badger population in Wytham Woods, Oxfordshire, UK (51^°^46:26 N, 1^°^19:19 W; for details see [42]) between 1995-2016, as part of an ongoing long-term research project. Following the methodology described in [56], all badgers received a permanent unique tattoo at first capture (typically as cubs: [42, 57], allowing individual identification (ID) and reliable aging.

Badgers were trapped in every month except during the closed season under the Protection of Badgers Act, 1992 (December-April), although, in some years, additional trapping was conducted under special license in December and early January before the end of the first pregnancy trimester [42]. The developmet of immature badgers could therefore be followed in: (1) *<u>spring</u>* (May/June) at end of the main mating period when cubs are fully weaned but spring weather can impact cub growth [58]; (2) *<u>summer</u>* (July/August/September) during additional mating activity previously reported in other high-density badger populations [29] and the period of lowest food abundance [46, 57]; (3) *<u>autumn</u>* (October/November) during reproductive quiescence and highest food abundance; and (4) *<u>winter</u>* (December/January) during the main mating season when cold weather may affect thermal energy balance in badgers [59].

### Somatic measurements and classification of external genitalia morphology

Head-body length (to the nearest 5mm), zygomatic arch width (to the nearest 1mm), and body weight (to the nearest 100g) were measured for all captured individuals, and a Body Condition Index (BCI) was calculated as log10(weight)/log10(body length). Subcaudal gland secretion was scooped out of the subcaudal pouch using a rounded stainless-steel spatula [32], and the volume estimated by eye to the nearest 0.05 ml. The spatula was disinfected between individuals using absolute ethanol [60]. External Genitalia Morphology (EGM) was recorded in both sexes and categorised according to Sugianto et al. [36] in females as normal, intermediate or swollen vulva, and in males as ascended, intermediate or descended testes. Male bacculum length, testes length, width and scrotal thickness were measured (in mm), and the testicular volume was calculated (in mm^3^) as (L x W x H) x 0.71, where L= testicle length – scrotal pinch, W= testicle width – scrotal pinch, and H = testicle width – scrotal pinch [61].

### Blood Sampling and Hormone Measurements

Blood samples (n_males_= 119; n_females_= 63; chosen from the available data set to represent each month – except the closed season, see above) were collected for endocrinological analyses via jugular venepuncture, using vaccutainer tubes (Becton-Dickinson) with K2-EDTA (ethylene diamine tetraacetic acid) anticoagulant. Sampling times were standardized to account for circadian variation in hormonal profiles [36-37], and blood samples were centrifuged within 30 minutes of sampling at 10°C for 10 min under 2,500 rpm/ 1470G. Plasma was transferred into Eppendorf tubes and frozen at ×20°C immediately.

All sex steroid titres were analysed using Enzyme-immunoassays (EIA) and analysed at the Chester Zoo Endocrinology Laboratory, UK. Oestrone was measured in microtitreplates coated with polyclonal antiserum raised against oestrone EC R522 [62]. Plasma samples were un-extracted and used for measurement after dilution with assay buffer at the ratio of 1:10. Duplicate 20µl aliquots of oestrone standard (0.195-200 pg/well), diluted plasma, and quality controls were combined with 50µl oestrone glucuronide coupled to horseradish peroxidase (oestrone-glucuronide-HRP) as label, and incubated at room temperature for 2 hours. Plates were washed five times and blotted dry after incubation, followed by an addition of 100 µL peroxidase substrate solution (ABTS) to each well. Plates were covered and incubated at room temperature until the ‘0’ wells reached approximately 1.0 optical density and read at 405 nm using a Spectrophotometer Opsys MR (Dynex). Assay sensitivity at 90% binding was 3.1 pg. Intra-assay coefficients of variation (CV, calculated as the average value from the individual CVs for all of sample duplicates), were 8.21 % (high) and 6.05 % (low); inter-assay variation (repeated measurements of high and low-value quality controls across plates) was 13.96 % (high) and 13.62 % (low) respectively.

Testosterone was measured in microtitre plates coated with anti-testosterone R156/7 (OEM-Concepts, UK). Samples (un-extracted) were analysed by dilution in 1:4 assay buffer. Duplicate 50µl aliquots of testosterone standards (2.3-600 pg/well), samples and quality controls were then combined with 50µl horseradish peroxidase (testosterone-HRP) as label. After incubation in the dark at room temperature for 2 hours, plates were washed 5 times and blotted dry, followed by addition of HRP-substrate (100 µL) to each well. Plates were covered and incubated at room temperature until the ‘0’ wells reached 1.0 optical density and were then read at 405 nm, using a Spectrophotometer (Opsys MR; Dynex). Assay sensitivity at 90% binding was 1.6 pg. The testosterone intra-assay coefficients of variation were 14.69 % (high) and 6.18 % (low), and inter-assay variation of high and low-value quality controls was 9.15 % (high) and 5.23 % (low).

## Statistical analysis

All statistical analyses were performed using RStudio (0.99.896) and R (R-3.2.4). Patterns of residuals, normality, and mean variance for each model were checked using R diagnostic plots. Generalized Additive Models (GAM) were used to generate trend lines for sex steroid levels (males: testosterone; females: oestrone) against age (3-28 months) using a smoothing function. A non-linear mixed model (random effect: badger identification/ tattoo number, ID) using the *nlme* and *sslogic* function was used to form a growth curve (providing an asymptote value as output) for bacculum length against age (3-28 months, n=773). To determine the age at which the bacculum ceased to grow, the percentage of the predicted bacculum length towards the asymptote (in the adult population) was calculated. Testes volume (n=597) and subcaudal secretion volume (n_male_=1233; n_female_=1284) trend lines were generated against age (3-28 months) by fitting a GAM model. Interactions between proportions of EGM (males: descended, intermediate, ascended testes, n=1136; females: normal, intermediate, swollen vulva, n=1174) with age (3-28 months) were analysed using a Chi-square test.

## Developmental heterochrony

### Endocrinology and EGM

Our GAM average trends (above) provided a legitimate basis of hormonal heterochrony in both sexes, where some cubs appeared to reach puberty earlier than others (i.e., the existence of two discrete groups that differ in their sex-steroid hormone levels at a certain age: high levels above and low levels below the GAM line benchmark, providing two developmental categories, or phenotypes). However, because endocrinological sample sizes were limited, we also repeated all analyses described below on EGM-based groups at the age of 11 months during their first mating season, which enhanced sample sizes. That is, in addition to comparing cubs with high vs low sex-steroid levels we also compared male cubs with ascended vs descended testes and female cubs with a normal vs swollen vulva; excluding intermediate conditions in both sexes to avoid ambiguity.

### Somatic growth

We subsequently compared head-body length, zygomatic arch width, BCI, and subcaudal secretion volume between these two groups using a linear model (including year as a factor to account for established inter-annual variation in growth patterns: [49, 52] to determine potential concurrent differences in physical development at the point of hormonal divergence. If significant differences were found in any of the skeletal size measures, the differences in head-body length and zygomatic arch width between adult size (above 28 months) to the size at the age at which divergence occurred were calculated in all individuals and compared between the respective groups, to determine heterochronous residual growth between the two groups. We then constructed growth curves for repeatedly captured individuals in each group based on the rates of increase in head-body length (which provide a reliable indicator for overall skeletal growth and development of badger cubs: [52]), employing a non-linear mixed model (random effect: ID) using the *nlme* and *sslogic* function. Where individuals were not recaptured at this precise target-age (within the 28 months period), we used the closest recapture point available for these analyses.

### Social factors affecting the timing of puberty

In addition, we investigated if social factors/ intra-specific competition affected the timing of sexual maturity by comparing hormone levels at the age when these phenotypic groups diverged with the total number of adults and cubs in that cub’s natal group and natal sett (as some groups utilize several setts: [63]), with year included as a factor. The total number of resident adults and cubs was determined annually by assigning residency according to the rules in Annavi et al. [50]; see also Sugianto et al. [52]).

## Results

### Endocrinological changes during the first 28 months

As predicted, throughout their first summer, all cubs had significantly lower sex-steroid levels than did adults. However, at the age of 11-12 months, i.e. during their first mating season, male cubs showed a small peak in testosterone (GAM: Edf= 8.631, R-sq.(adj)= 0.566, GCV= 2.230, Deviance explained= 59.9%, p<0.001, Fig 1), which was then followed by the seasonal pattern of testosterone levels typical for adult males (high in spring and summer, and low in autumn: [37]), with a pronounced peak that reached levels typical for adults during their second mating season (22-28 months).

**Figure 1.**
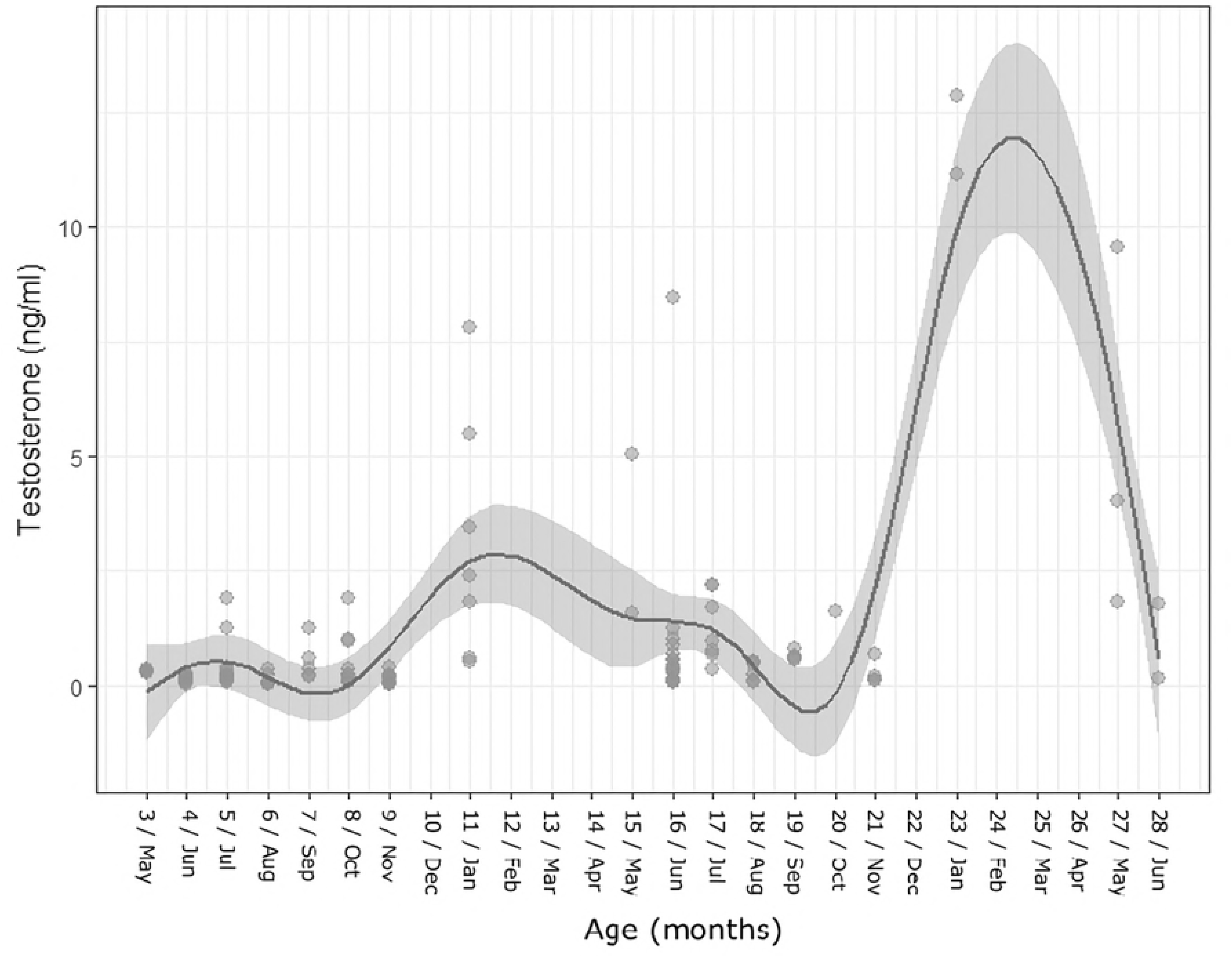
Testosterone levels (ng/ml) in males aged 3-28 months.

In females, oestrone levels increased gradually from the age of 3 up to 11 months, at which point they almost reached adult levels (GAM: Edf= 5.218, R-sq.(adj)= 0.216, GCV= 733.1, Deviance explained= 28.2%, p=0.009, Fig 2). Patterns then followed the seasonal oestrone pattern typical for adults, with levels being relatively high in spring (13-16 months) and decreasing in summer (17-19 months), remaining low during autumn (20-21 months, reproductive quiescence) and winter (22 months, December: implantation) after which time they increased again towards spring (27-28 months), this time reaching adult levels (73.28±28.06 pg/ml; [30]). Nevertheless, inter-individual variation among females was considerable during the second summer (months 15-20), i.e., from the end of their first mating season until autumnal reproductive quiescence.

**Figure 2.**
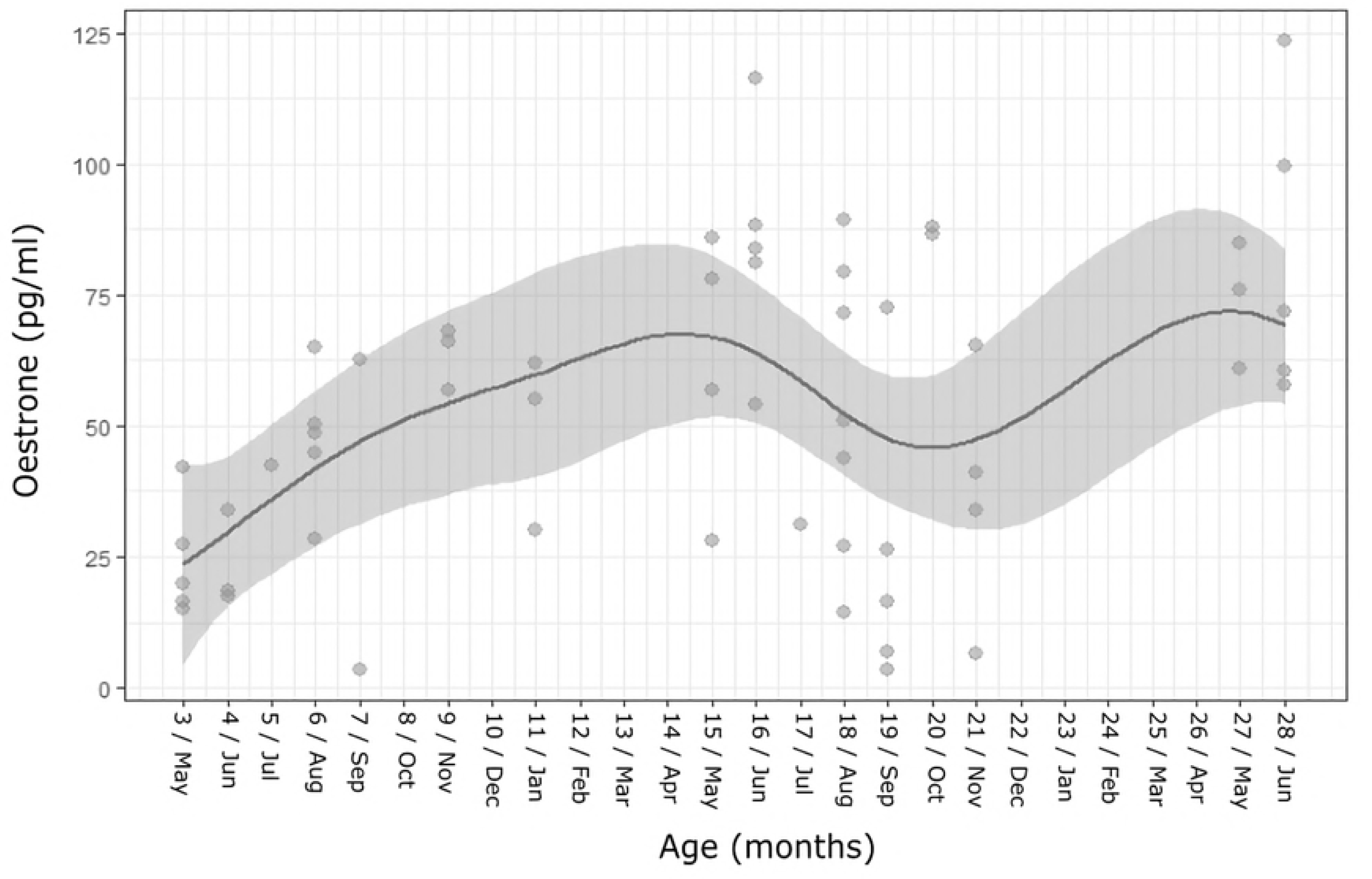
Oestrone levels (pg/ml) in females aged 3-28 months.

### Changes in external genitalia during the first 28 months

In both sexes, there was a significant interaction between EGM and age (in months) (males_n_=1136: X^2^= 937.04, df = 36, p<0.001; females_n_=1174: X^2^= 418.47, df = 34, p<0.001; Fig 3a, b). The majority of male cubs had scrotal (i.e., fully descended) testes for the first time at the age of 5-6 months (83.3% and 70.4% respectively), while during their first autumn (8-9 months) the majority had ascended testes (63.6% and 77.1% respectively). During their first mating season in January (11 months), the largest proportion (41.5%) of male cubs had descended testes, while both ascended and intermediate proportions were each 29.3%. During the following spring (15 months) and summer (19 months) the majority of males (94.8% and 82.9%, respectively), had descended testes and followed the adult seasonal pattern thereafter.

**Figure 3.**
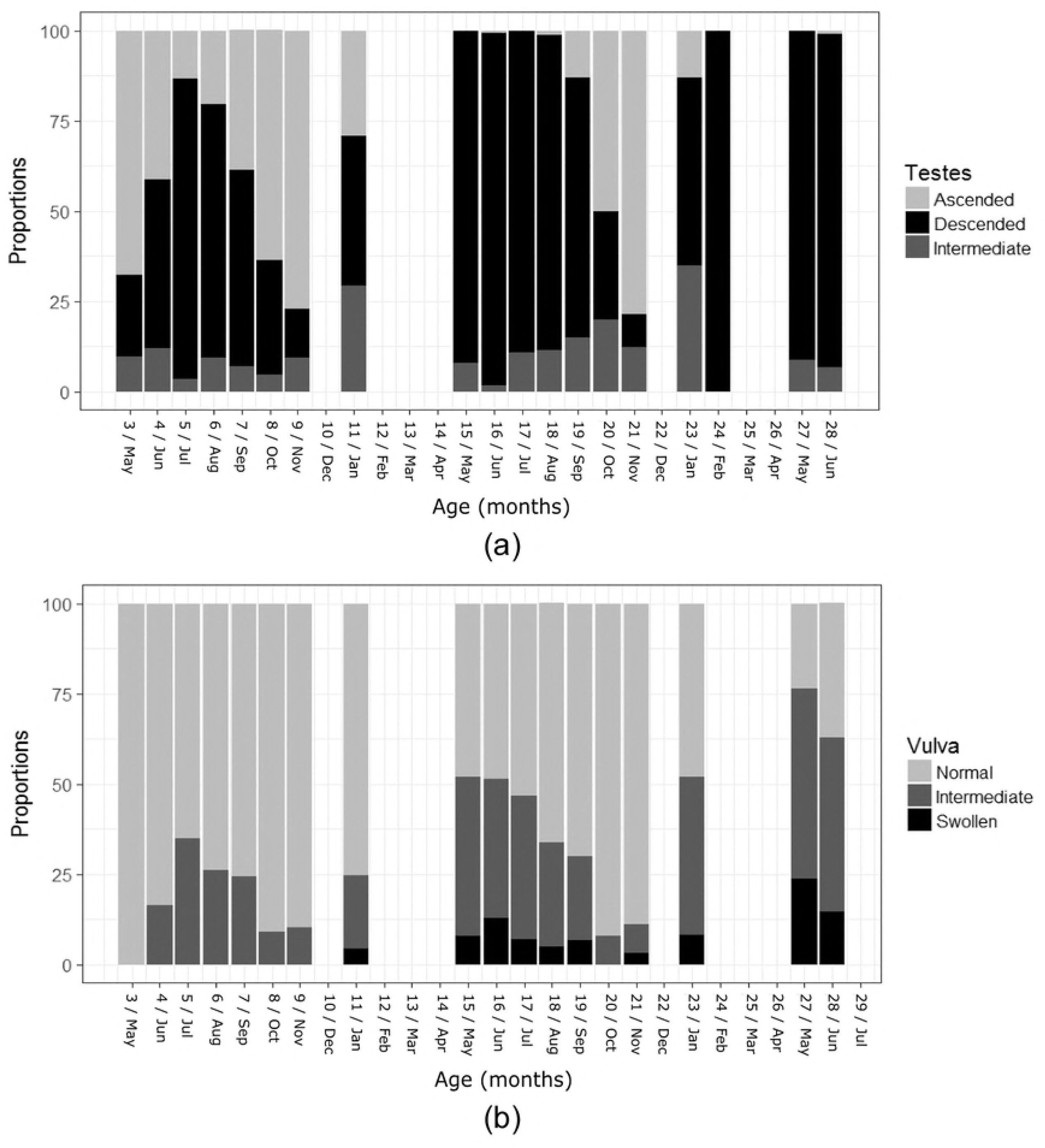
EGM changes in males (a) and females (b) aged 3-28 months.

In females, the earliest vulval swellings (4.4% swollen; 20% intermediate vulva) were recorded at the age of 11 months during their first mating season (Jan). The proportions of females with intermediate and swollen vulva increased during the next spring-summer (15-19 months; intermediate: spring= 41.4%, summer= 30.5%; swollen: spring= 10.3%, summer= 6.2%) and decreased in autumn (20-21 months; intermediate: 7.8%, swollen: 1.5%). During their third spring, the highest percentage of female cubs had either intermediate (50.4%) or fully developed vulval swelling (19.2 %), congruent with adult states [36].

In males, testicular volume started to increase markedly at the age of 11 months (from 848.69±475.20 mm^3^at the age of 4-6 monthsn=12; 1331.27±1289.97 mm^3^ at the age of 7-9 month n=32; to an average of 3449.73±1572.04 mm^3^ at 11 months n=18) and peaked during the first mating season (5326.45±2674.88 mm^3^, n=60; winter-early spring; GAM: Edf= 6.921, R-sq.(adj) = 0.242, Deviance explained = 25.1%, p>0.001, Fig 4). Average testicular volume then decreased towards an autumnal minimum at 20-21 months (3209.42±2283.41 mm^3^, n=20) and followed the adult seasonal pattern thereafter, reaching a slightly higher peak (5698.26±2409.85 mm^3^, n=187) in the second mating season (22-28 months), with sizes comparable to adult values (winter: 6650.82 mm^3^, spring: 5776.31 mm^3^: [36]).

**Figure 4.**
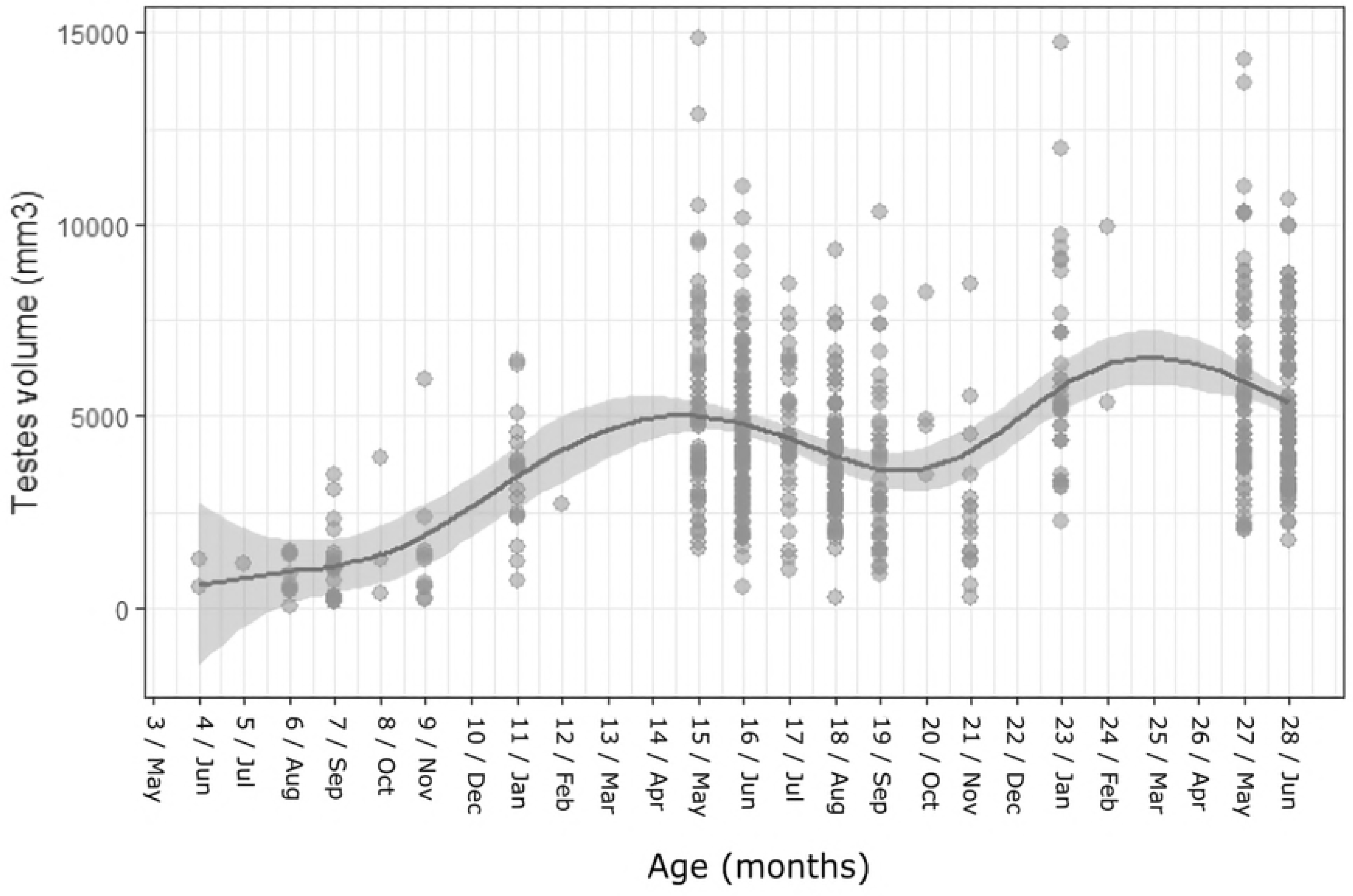
Testes volume (mm^3^) in males aged 3-28 months.

Male bacculum length increased consistently month by month for the first year, when bacculum growth rates slowed and reached 99% towards the asymptote of 86.03 mm predicted by our model at the age of 23-24 months (Fig 5).

**Figure 5.**
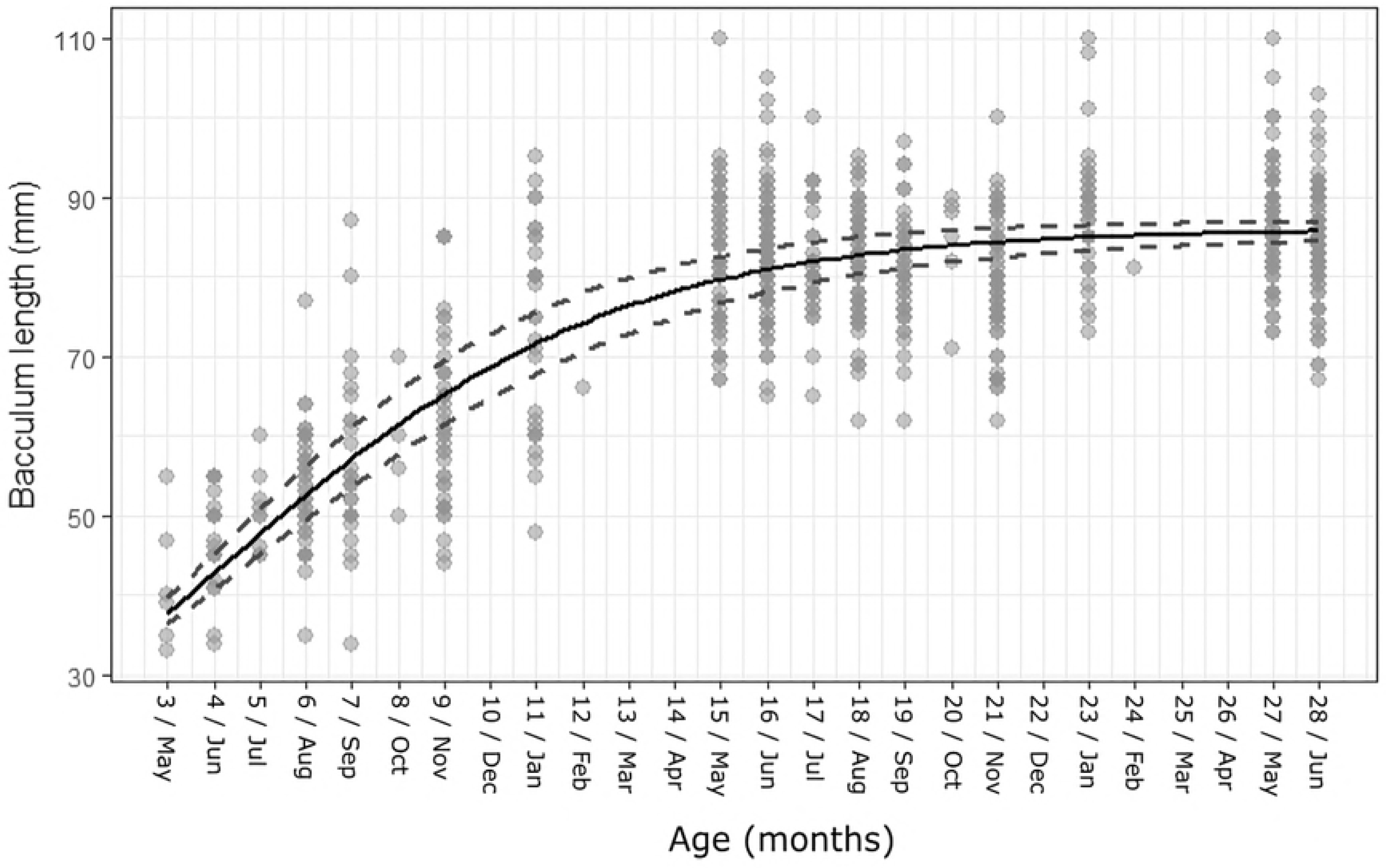
Bacculum length growth curve at age of 3-28 months in males.

### Subcaudal gland activity during the first 28 months

Both sexes started producing subcaudal gland secretion at a similar age (during first capture at 3 months; Fig 6). Nevertheless, secretion volume was very low (unmeasurable traces in males and 0.04±0.09 ml in females _n=72_) and increased only slowly towards their first mating season at age 11 months (0.38±0.36 ml in males_n=41_and 0.19±0.13 ml in females_n=47_). Thereafter, secretion volume increased substantially in both sexes (GAM: male: Edf= 8.45, R-sq.(adj)= 0.56, GCV= 0.18, Deviance explained= 56.2%, p<0.001, Fig 6a; female: Edf= 8.83, R-sq.(adj)= 0.37, GCV= 0.04, Deviance explained= 38.2%, p<0.001, Fig 6b). In male yearlings, secretion volume peaked in spring-summer (13-18 months, 1.04±0.61 ml, n=292), decreased towards an autumn-minimum (20-21 months; 0.56±0.39 ml, n=97) and peaked again (with higher secretion volume) in their second winter-spring (23-28 months; 1.17±0.58 ml, n=210), following the typical seasonal pattern and secretion volume of adults (average values for spring= 1.06±0.67 ml, n=1004; summer= 0.92±0.66 ml, n=1059; autumn= 0.60±0.50 ml, n=790; winter= 0.90±0.60 ml, n=347 ml). Female yearlings showed a first slight peak in secretion their second spring (13-16 months; 0.35±0.24 ml, n=150), after which volume decreased slightly during summer-autumn (17-19 months, 0.32±0.35 ml, n=189 and 20-21 months, 0.27±0.17 ml, n=110), but then started to increase again at the end of winter (24 months), peaking at an average of 0.52±0.41 ml in spring (27-28 months), and following the adult pattern thereafter (average values for spring = 0.41±0.34 ml, n = 1287; summer = 0.33±0.32 ml, n = 1286; autumn = 0.25±0.24 ml, n = 933 ml; winter = 0.24±0.26 ml, n = 207).

**Figure 6.**
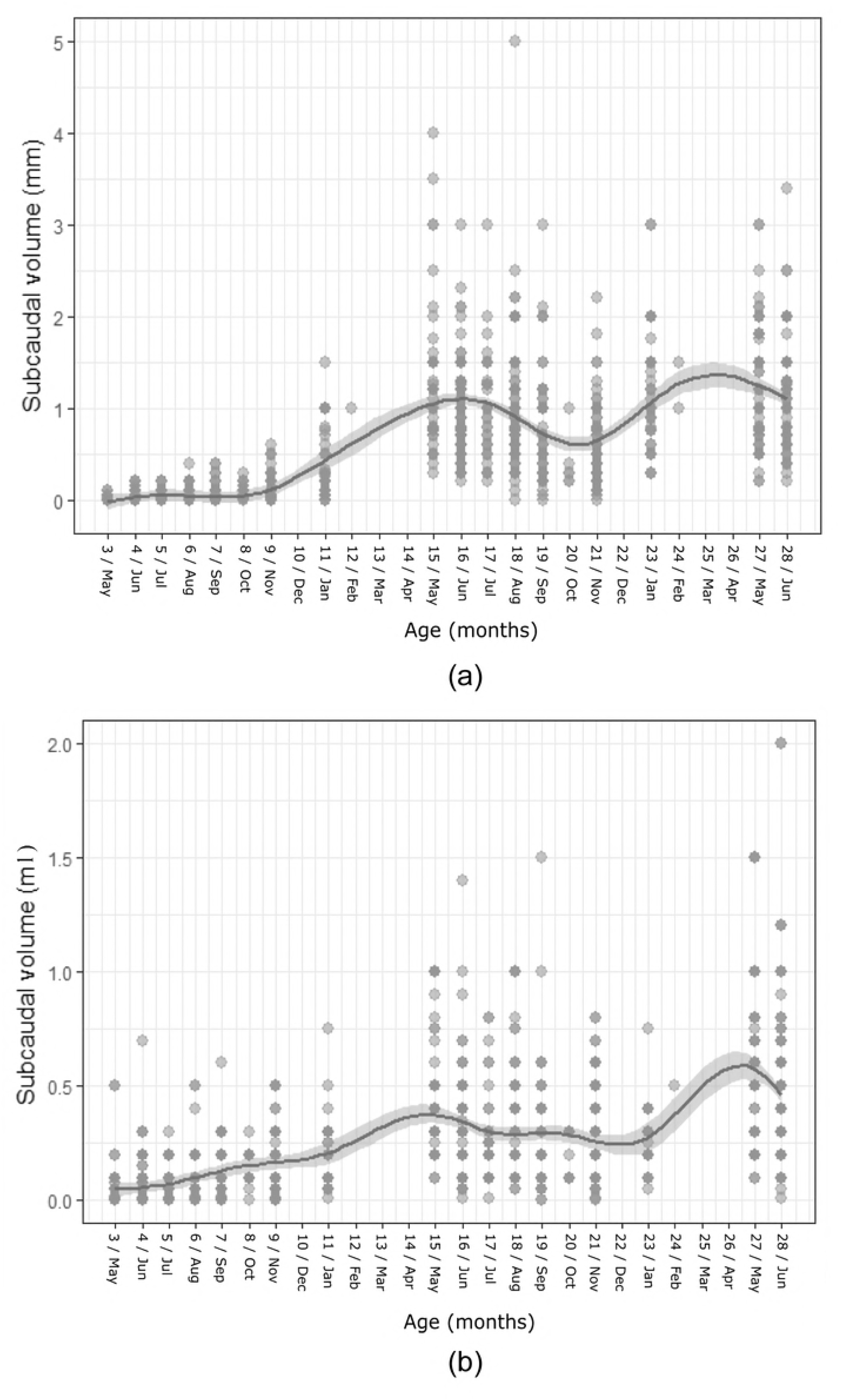
Subcaudal secretion volume (ml) changes in males (a) and females (b) aged 3-28 months.

### Evidence for two categories of cubs: Early-and Late-Developers

#### Endocrinological evidence for early and late developers

In both sexes, some individuals appeared to reach puberty earlier than others evidencing the existence of two endocrinological phenotypes: early-and late developers (Fig 1, 2), clustering into two distinct trait types according to the GAM line benchmark which exposed significantly different sex-steroid levels (for detailed results see Table 1 and 2). However, the age at which these early-and late development categories became apparent, differed between male and female cubs. For males, some (3/7= 42.9%) cubs reached puberty during their first year (HT, testosterone levels above the GAM line at 11 months of age), while the remainder reached puberty during their second year (LT, testosterone levels below the GAM line at 11 months, reaching pubescent levels at 22-28 months of age; Fig 1). In contrast, in females, the two possible endocrinological phenotypes diverged at age 15 -18 months (younger cubs either had more unified levels or sample sizes were too small to signify a difference; Fig 2), where some females had above-average oestrone levels (HO; oestrone above the GAM line) and some below-average levels (LO; oestrone below GAM line). In both sexes, these endocrinological phenotypes manifested independent of calendar year.

**Table 1.**
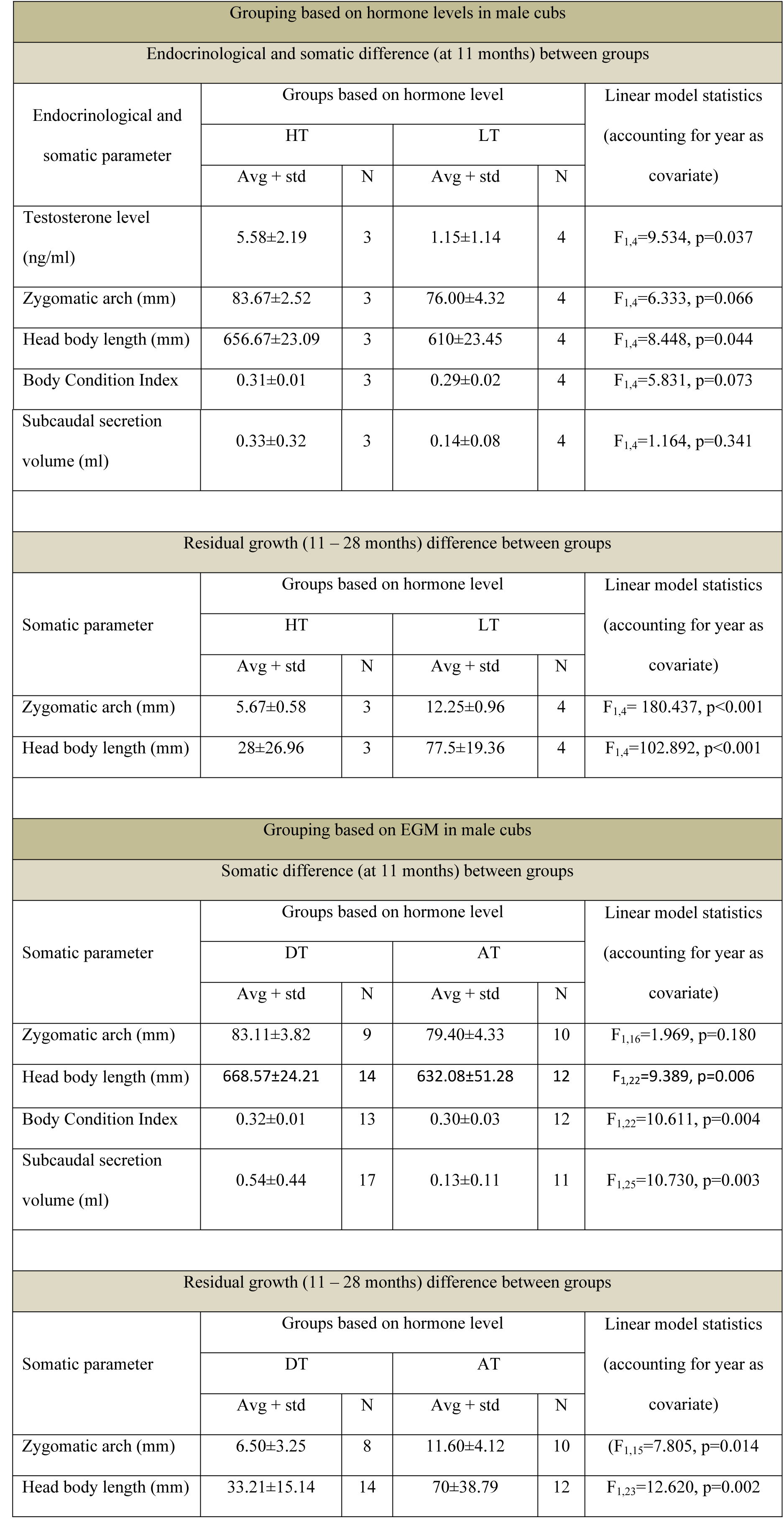
Differences in somatic development between early and late developers from endocrinological and EGM grouping in male cubs.

**Table 2.** Differences in somatic development between assumed phenotypes from endocrinological and EGM grouping in female cubs.

| Grouping based on hormone levels in female cubs |  |  |  |  |  |
| --- | --- | --- | --- | --- | --- |
| Endocrinological and somatic difference (at 15 - 18 months) between groups |  |  |  |  |  |
| Endocrinological and somatic parameter | Groups based on hormone level |  |  |  | Linear model statistics<br>(accounting for year as covariate) |
|  | HO |  | LO |  |  |
|  | Avg + std | N | Avg + std | N |  |
| Oestrone level (ng/ml) | 86.06±12.72 | 9 | 38.31±15.32 | 8 | F<sub>1,14</sub>= 48.283, p<0.001 |
| Zygomatic arch (mm) | 80.44±3.81 | 9 | 81.29±3.15 | 7 | F<sub>1,13</sub>= 0.219, p=0.647 |
| Head body length (mm) | 672.22±23.6 | 9 | 662.5±35.15 | 8 | F<sub>1,14</sub>= 0.700, p=0.417 |
| Body Condition Index | 0.3±0.02 | 9 | 0.3±0.04 | 8 | F<sub>1,14</sub>=0.014, p=0.909 |
| Subcaudal secretion volume (ml) | 0.48±0.42 | 8 | 0.9±1.46 | 5 | F<sub>1,10</sub>= 0.273, p=0.613 |
| Grouping based on EGM in female cubs |  |  |  |  |  |
| Somatic difference (at 15 - 18 months) between groups |  |  |  |  |  |
| Somatic parameter | Groups based on hormone level |  |  |  | Linear model statistics<br>(accounting for year as covariate) |
|  | SV |  | NV |  |  |
|  | Avg + std | N | Avg + std | N |  |
| Zygomatic arch (mm) | 83.41±2.22 | 22 | 82.85±3.96 | 124 | F<sub>1,143</sub>= 0.446, p=0.505 |
| Head body length (mm) | 678.1±18.54 | 21 | 675.53±26.59 | 148 | F<sub>1,166</sub>= 0.188, p=0.665 |
| Body Condition Index | 0.3±0.01 | 21 | 0.3±0.02 | 147 | F<sub>1,165</sub>= 0.051, p=0.822 |
| Subcaudal secretion volume (ml) | 0.36±0.22 | 21 | 0.32±0.21? | 142 | F<sub>1,160</sub>=0.574, p=0.450 |
| Somatic difference (at 11 months) between groups |  |  |  |  |  |
| Somatic parameter | Groups based on hormone level |  |  |  | Linear model statistics<br>(accounting for year as covariate) |
|  | SV |  | NV |  |  |
|  | Avg + std | N | Avg + std | N |  |
| Zygomatic arch (mm) | 83.5±7.78 | 2 | 78.2±3.79 | 25 | $F_{1,24} = 3.105, p=0.091$ |
| Head body length (mm) | 675±21.21 | 2 | 645.88±37.43 | 34 | $F_{1,33} = 1.247, p= 0.272$ |
| Body Condition Index | 0.32±0.02 | 2 | 0.31±0.03 | 34 | $F_{1,33} = 0.764, p=0.388$ |
| Subcaudal secretion volume (ml) | 0.2±0.0 | 2 | 0.18±0.14 | 33 | $F_{1,33} = 0.221, p=0.641$ |

### Differences in somatic development between early and late developers

**Males**

Comparing the extent of somatic development between the two endocrinological phenotypes (at age 11 months), we found that HT males (n=3) had significantly larger head-body length than LT cubs (n=4), were larger overall, and showed a near significant difference in zygomatic arch width and BCI; but did not differ in the volume of subcaudal gland secretion they produced.

To test whether the early (endocrinological) development of individuals assigned to the HT group was simply the product of a more rapid development to adult size and were not just larger cubs but indeed early-developers (and were thus closer to being fully developed, sexually mature adults), we compared the differences in head-body length and zygomatic arch width at the age above 28 months (i.e., fully developed adults) with those of cubs assigned to HT-and LT endocrinological categories at the age of 11 months. The difference in these measurements between HT-cubs and adults was significantly smaller than in LT-cubs and adults, confirming that HT-cubs were closer to being fully developed adults (see Table 1).

For males, comparing the growth curves of HT (8 repeat-measures over 33 months from 3 individuals) and LT types (32 repeat-measures over 33 months from 4 individuals) revealed a trend (albeit non-significant, likely due to limited sample sizes) for LT-cubs to grow more slowly than HT-cubs, despite ultimately reaching similar adult head-body lengths (X^2^=1.873, df=7, p=0.392; non-linear mixed model), where HT had already reached 95% of the maximum head body length at the age of 11 months, whilst LT reached this percentage later at the age of 14 months. This difference in body size disappeared at the age of 19-20 months, when growth rates of HT-and LT-males equalised (Fig 7).

**Figure 7.**
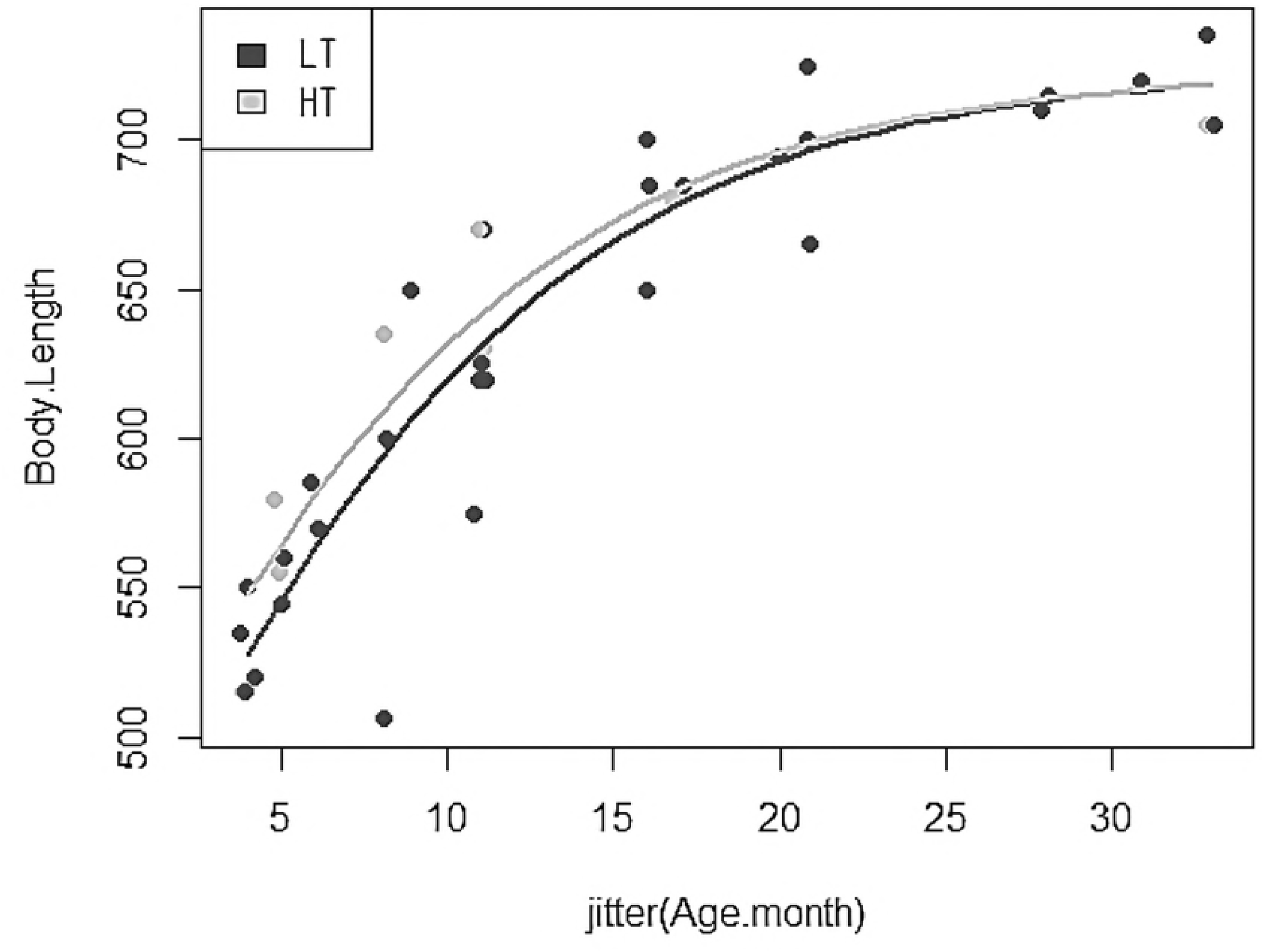
Body length growth curve of HT (open circles) and LT (solid black circles) groups, age: 4-33 months.

We repeated these analyses based on the degree of testicular descent at the age of 11 months, comparing cubs that had fully descended testes (DT; assumed to have reached puberty; n=18) with those that had ascended testes (AT; assumed to have not reached puberty; n=13). Overall, these showed similar differences in somatic development to categories arising according to endocrinological groups (for detailed results see Table 1). DT cubs were considerably longer (head-body length), were in significantly better body condition/BCI, and had significantly more subcaudal secretion at 11 months of age than AT males; but no difference was found in zygomatic arch width. DT cubs also had significantly smaller adult-cub differences in head-body length and zygomatic arch width than AT cubs, which corroborated that (like HT-males) they were closer to adulthood.

Mirroring the trend found in the endocrinological phenotypes, there was a significant difference in the growth curve of these two groups (Fig 8; X^2^=10.087, df=8, p=0.018), with DT cubs (117 repeat-measures taken over 35 months from 18 cubs) growing faster, and reaching adult size earlier than AT cubs (99 repeat-measures taken over 35 months from 13 cubs). At the age of 19-20 months this difference disappeared and growth rates of DT and AT cubs equalised.

**Figure 8.**
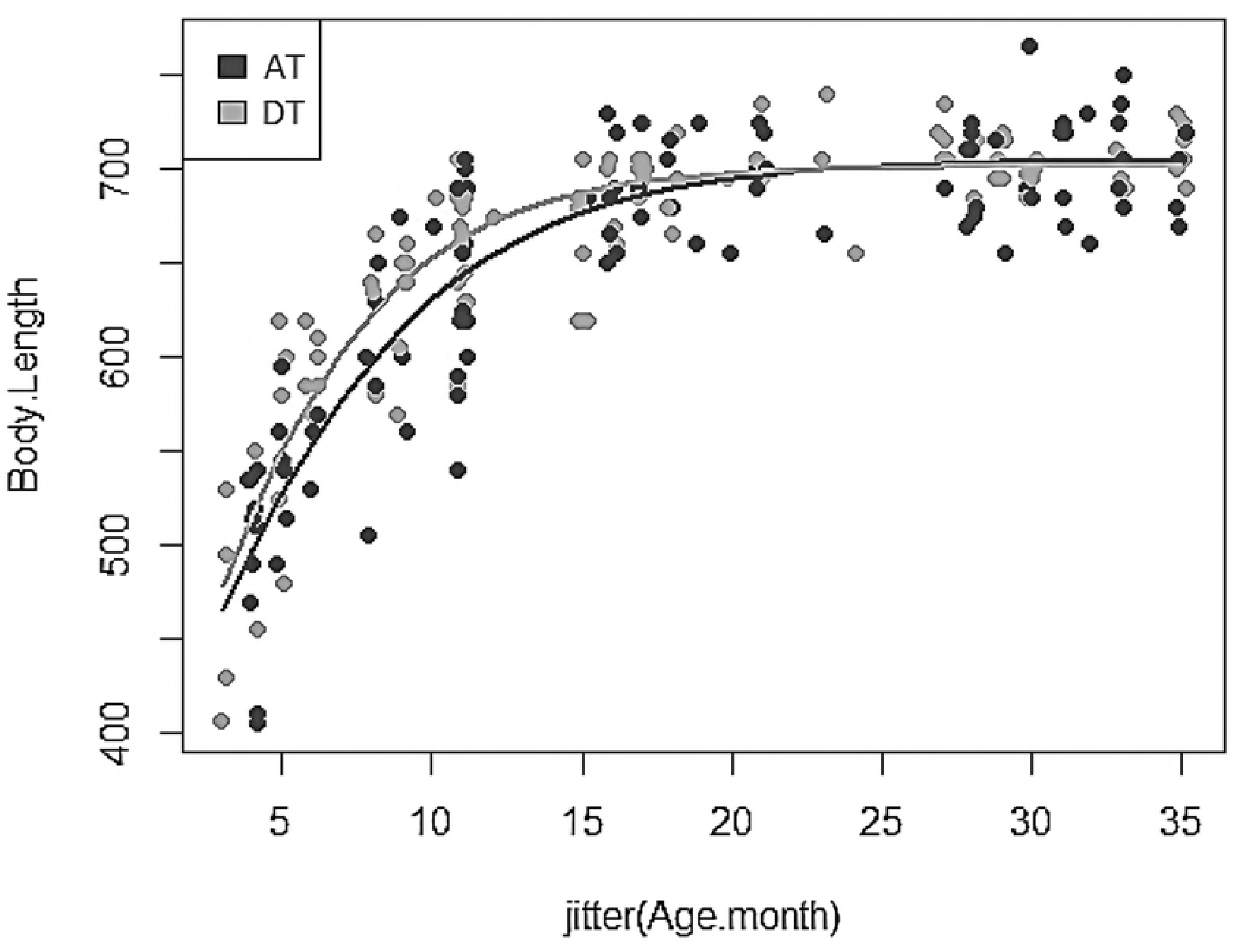
Body length growth curve of descended testes (DT) cubs and ascended testes (AT) cubs, age: 3-35 months.

### Females

In contrast, for females, we detected no (significant) differences between any of the somatic parameters, nor for subcaudal gland volume, when comparing between the early (nHO= 9) and late (nLO= 8) developing endocrinological phenotypes (see Table 2). To ensure that the somatic similarity found between the two endocrinological phenotypes was not an artefact of the smaller sample size for our endocrinological dataset, we used vulva category at the ages between 15-18 months as the criteria defining stage of development. Again, on this basis, we also found no significant differences (see Table 2) in any somatic parameters nor subcaudal gland volume between SV cubs (n= 24) and NV (n=152), evidencing that both vulva condition types exhibited similar body size by age.

However, because some female cubs first exhibit vulval swelling at the age of 11 months (n_SV_=2, n_NV_=34), we repeated these analyses (see Table 2) at this younger age, but again found no significant difference in BCI, head-body length, nor subcaudal gland volume, although we did detect a slight difference in zygomatic arch between these groups.

### Social factors influencing the timing of puberty

At the age of 11 months (see Table 3), testosterone levels in male cubs tended to be lower in larger natal social groups and setts, albeit without statistical significance, although the high R-value (0.53 and 0.63 for resident adults in natal social group and sett respectively) evidences a strong correlation, where non-significance is likely due to low sample sizes; with a similar interaction evidenced by an even higher R-value with number of other cubs present in the natal social group and sett (0.76 and 0.85 for other resident cubs in natal social group and sett repectively). That is, cubs born/ growing up in smaller social groups and/ or setts may be more likely to be early developers than those born/ growing up in larger groups/ setts. Using degree of testicular descent at the age of 11 months instead of the endocrinological phenotype, however, suggested no trend (see Table 3).

**Table 3.** Social factors affecting the timing of puberty in male and female cubs.

| Social factors affecting timing of puberty in male cubs (11 months) |  |  |
| --- | --- | --- |
| Social factor | Testosterone | EGM |
| Number of adults in natal social group | R= -0.53; $F_{1,4}= 1.407$, p=0.301 | R= -0.39; $F_{1,26}= 1.052$, p=0.314 |
| Number of adults in natal sett | R= -0.631; $F_{1,3}= 1.446$, p=0.316 | R= 0.31; $F_{1,23}= 1.994$, p=0.171 |
| Number of cubs in natal social group | R= -0.763; $F_{1,3}= 3.527$, p=0.157 | R= 0.36; $F_{1,22}= 2.543$, p=0.125 |
| Number of cubs in natal sett | R= -0.851; $F_{1,4}= 5.359$, p=0.082 | R= 0.46; $F_{1,17}= 2.555$, p=0.128 |
| Social factors affecting timing of puberty in female cubs (15 – 18 months) |  |  |
| Social factor | Oestrone | EGM |
| Number of adults in natal social group | R= -0.157; $F_{1,14}=0.0393$, p=0.846 | R= -0.1; $F_{1,172}=0.272$, p=0.603 |
| Number of adults in natal sett | R= -0.116; $F_{1,14}=0.016$, p=0.901 | R= -0.23; $F_{1,165}=2.281$, p=0.133 |
| Number of cubs in natal social group | R= -0.267; $F_{1,14}=0.4299$, p=0.523 | R= 0.02; $F_{1,144}=0.002$, p=0.963 |
| Number of cubs in natal sett | R= -0.092; $F_{1,13}=0.052$, p=0.823 | R= -0.02; $F_{1,102}=0.005$, p=0.944 |

Female cubs showed only very weak negative relationships between oestrone levels at the age of 15-18 months with the number of adults resident in their natal social group and sett, as well as the number of other cubs in their natal social group and sett. When cubs were categorised on the basis of their EGM at the age of 15-18 months, again no effects were found (see Table 3).

## Discussion

We demonstrate that, in badgers, puberty begins in both sexes at ca. 11 months of age, when cubs develop similar seasonal sex-steroid patterns to adults. Furthermore, in both sexes, all parameters that support reproductive activity and mating-associated behaviours, such as external genitalia morphology (and in males also testes volume), and subcaudal gland secretion volume, show similar developmental patterns to sex steroid levels, further corroborating the onset of puberty [1]. The increase of sex-steroid levels likely triggers changes in EGM [36], as well as in the activity of species-specific subcaudal glands important in the context of reproduction and sexual advertisement [64]. Nevertheless, cub titres typically remained lower than those reported for adults (males: [37]; females: [30]) until their second mating season (22-26 months).

In male cubs, testosterone levels remained low and exhibited no seasonal variation until the first winter mating season, when they started to increase, reaching a smaller peak than in adults [37] Levels then remained elevated until the end of the mating season, and followed the seasonal pattern typifying adults thereafter. By the time male cubs reached the second population breeding season, their testosterone titres had reached higher levels (compared to the first mating season) in accord with adults [37]. Bacculum growth also reached the population-based asymptote by the cubs’ second mating season, indicating the completion of sexual development. These findings support the hypothesis by Whelton and Power [38], who measured bacculum length post-mortem in road kill and culled badgers, positing that the observed abrupt decrease in bacculum growth rate coincides with sexual maturity; although their post-mortem study was unable to verify this conclusion through endocrinological measurements.

In contrast, in females, we observed a gradual increase in oestrone from the age of 3-11 months (May-January) without any noticeable seasonal variation. After age 11 months, however, cub oestrone levels started to follow the same seasonal pattern as adults [30], with high levels in spring and low levels in summer. Nevertheless, although adult females oestrone levels typically increase in autumn and remain elevated until blastocyst implantation in December [30], cub oestrone levels remained low until the next mating season (January), implying that – counter to observations from other, low-density studies [54] -no female cubs in our dataset were capable of mating successfully during their first year, corroborating genetic results from our study population reported previously [65].

Inter-individual variation in plasma sex-steroid levels, however, was considerable among same-age cubs of either sex, and we observed two distinct categories: early and late developers. We infer these to qualify as distinct phenotyptic response types, given the potential fitness advantage of early maturity [66], but set against the reality of resource limitation and social stress in wild populations typically precluding all individuals from engaging in the maximal developmental response [51]. In male cubs, endocrinological profiles and EGM indicated that some had reached sexual maturity at the age of 11 months, while the remaining cubs likely achieved this only during their second winter. At this time all male cubs showed similarly large testosterone peaks comparable to adult levels and puberty had concluded. Similarly, there was substantial variation in the proportion of the final body length males had achieved by age 11 months, mirroring the differences in testosterone levels observed during this period. This increase in head body length ceased by age ca. 18 months (99 % towards asymptote; [52]) which is also the age at which body lengths equalised when dividing males according to both testes descent categories (DT and AT as well as endocrinological categories HT and LT). Nevertheless, when we cross-referenced against assigned paternity data, none of the individuals exhibiting high hormone levels at 11 months (HT=3 individuals) were assigned cubs the spring after. For female cubs, variation in oestrogen levels was high during their second summer (May-Sept/ Oct), indicating that not all females reached sexual maturity at the same age, but that puberty onset varied between 15-18 months of age.

The existence of different ontological phenotypes (i.e., early and late developers) has been described in other members of the Mustelidae, and has been linked to body size and species-specific life history traits [27] For example, in captive sables (*Martes zibellina*), a small proportion of all individuals are reported to reproduce at 15 months of age, whereas the majority (80%) of males and females starts breeding at the age of 27 months [27]. Our results support observations from badgers in Sweden, where spermatozoa were first recorded in male cubs at the age of 12 months [54]. Nevertheless, in this Swedish low-density population, the majority of males reached puberty in their first year, and only a minority of males did not produce spermatozoa until their second summer, or even winter (24 months: [54]). In our high-density population, in contrast, most males reached puberty in their second year.

Overall, our results support the hypothesis that mammals typically need to reach a threshold body size for sexual maturation [10]. Thus, the age at which puberty occurs is likely not only influenced by the gene load of the individual but also by ecological factors such as access to food (affected by weather/ climate), competition, and differences in demographic factors [1]. Resource availability tends to vary across time and space [67], and access to resources is further constrained by the number of competing conspecifics present leading to social stress [63]. Consequently, energy budgets can differ substantially between individuals even within a single population/ year, with the potential to drive considerable variability in the timing of sexual maturity [1]: Individuals that develop under poor nutritional conditions, or subject to more social stress resulting from competition, usually reach sexual maturity at slower rates [11-13] as implied by our observation that male cubs born into bigger setts and larger social groups tended to be biased toward the late developer phenotype.

Generally, in mammals (especially those with polygynous mating systems), males tend to grow more quickly than females [68], and ultimately attain a larger body-size (i.e., dimorphism; see Badyaev [69]). Our data show that this is also the case in badgers (see also Sugianto et al. [52]; NB, our measurements were made after weaning, and thus obviating differential maternal investment effects; [70]). Thus, males are likely also more vulnerable to resource limitation and social competition, with the potential to impact their degree of development by the end of their first year, explaining the observed delay in puberty in larger social groups [71-72]. Our findings, are congruent with those for female brown bears (*Ursus arctos*), where adult body size shows a negative relationship with population density [73]: female bears are larger and grow faster in areas with better environmental conditions, while with higher resource competition, females are smaller and grow more slowly. As in our badger study, bears have been shown to compensate for slower growth rates by delaying reproductive activity [73] at the potential cost of lower lifetime reproductive success [74]. Similar negative correlations between population density and individual growth rate has been reported in the northern fur seal (*Callorhinus ursinus*; [75]), polar bears (*Ursus maritimus*; in terms of smaller juvenile body length [76], and adult body size [77]), and in American black bears (*Ursus americanus*; with lower yearling weight; [78]). Similarly, in farmed red deer (*Cervus elaphus*), [79] found that the growth rate of subordinate females was 2.5 times slower, and average daily weight gain of all juvenile hinds was significantly impaired, under high stocking density. Demographic effects have also proven to affect the onset of maturation in female baboons (*Papio cynocephalus*), where first mentruation was earlier in smaller groups where individuals experience less social stress and competition [80].

We thus conclude that the asynchronous timing of puberty, leading to two heterochronous phenotypes, can occur even within a single population, and is likely caused by individuals attaining the required minimum body size according to different time scales. Ultimately, capacity to breed at a young(er) age can have profound effects on life-history trade-offs (see [81] with early-life success often being critical to an individual’s fitness [66] and can substantially enhance population growth rate [82].

## Acknowledgements

We acknowledge the long-term support of the People’s Trust for Endangered Species (PTES) for the Wytham Badger Project. NAS was supported by a DPhil scholarship from Indonesia Endowment for Education 2014-2018, and CDB was supported by a Research Fellowship from the Poleberry Foundation. Dr. Sue Walker’s help for hormone analyses in Chester Zoo Laboratory, Chester is warmly acknowledged. All protocols and procedures employed were approved by the Animal Welfare and Ethical Review board of Oxford University’s Zoology Department and procedures were conducted under the Animals (Scientific Procedures) Act, 1986 and Natural England licenses (PPL: 30/3379)

